# Topical heterogeneity in affective touch: Does it impact body image?

**DOI:** 10.1101/2020.11.30.403600

**Authors:** Valentina Cazzato, Sofia Sacchetti, Shelby Shin, Adarsh Makdani, Paula Diane Trotter, Francis McGlone

**Author notes:** **Correspondence to:** Dr Valentina Cazzato, School of Psychology, Faculty of Health, Liverpool John Moores University, UK.

## Abstract

Recent evidence suggests that altered responses to affective touch - a pleasant interoceptive stimulus associated with activation of the C-Tactile (CT) system, may contribute to the aetiology and maintenance of mental conditions characterised by body image disturbances (e.g., Anorexia Nervosa). Here, we investigated whether tactile pleasantness and intensity differ across body sites, and if individual differences in dysmorphic appearance concerns and body awareness might be associated to touch perceptions across body sites. To this end, we measured perceived pleasantness and intensity of gentle, dynamic stroking touches applied to the palm, forearm, face, abdomen and back of 30 female participants (mean age: 25.87±1.17yrs) using CT-optimal (3 cm/s) and non-CT optimal (0.3 and 30 cm/s) stroking touch. As expected, participants rated CT-targeted touch as more pleasant compared to the two non-CT optimal stroking touch at all body sites. Nevertheless, CT-targeted touch applied to the abdomen elicited the lowest pleasantness ratings compared to all other body sites and to the two non-CT optimal stroking touch. Individual differences in body awareness and dysmorphic concerns significantly predicted preference for CT-optimal over non-CT optimal stroking touch applied to the forearm and the back. These findings begin to elucidate the link between CT sensitivity, dysmorphic appearance concerns and body awareness, which may have implications for future research looking to inform early interventions. Addressing impaired processing of affective interoceptive stimuli, such as CT-targeted touch, may be the key to current treatment approaches available for those populations at risk of disorders characterised by body image disturbance.

## Introduction

Touch is a crucial means of receiving information from the outside world by mediating our interactions with objects, and other individuals. The touch experience is a combination of the discriminative aspects of tactile perception (i.e. characterising and localising external stimuli), and the affective and social qualities encoded therein [1, 2]. Previous investigations have demonstrated that slow, caress-like touch is usually experienced as highly pleasant [3–6], and plays a pivotal role in instigating and maintaining close social interactions.

It is hypothesised that social and affective qualities of touch are mediated by a specific class of unmyelinated mechanoreceptors – C-tactile afferents (CTs) [7–9], typically found in hairy but not glabrous skin. Neurophysiological investigations by means of single-unit microneurography have shown that CTs are velocity tuned, responding optimally to low force stroking velocities (~ 3cm/s), delivered at skin temperature. The relationship between CT activation and stroke velocity is best described by an inverted U-shaped regression, with the greatest response at 3 cm/s, and weaker responses at slower (0.1 cm/s) and faster velocities (30 cm/s) [3, 4]. Importantly, this response pattern strongly correlates with subjective ratings of stimulus pleasantness and velocity preference [3].

According to the ‘Social Touch hypothesis’ [1] CTs, which carry information about the ‘hedonic’ aspect of interpersonal, gentle touch, have evolved to promote social bonding. A multitude of research has corroborated their importance in affiliative interactions. It has been shown that, when asked to stroke their partners or babies, people spontaneously delivered touch at CT-optimal velocities [10]. Moreover, CTs optimally respond to touch delivered at skin temperature [5]. Consistent with the social touch hypothesis, CT-optimal touch was also found to have physiological, soothing effects [11–13], and health benefits, including stress relief associated with a decrease in heart rate, blood pressure, and cortisol release [14]. For example, recent psychophysiological investigations have shown that CT-optimal stroking to the forearm of 9-month old infants selectively decreased heart rate [13] and that CT-targeted touch produces a significant decrease in preterm infants’ heart-rates and increase in their blood oxygenation levels [14]. Furthermore, a recent study [15] reported that affective touch reduced the distress caused by social exclusion.

The primary cortical target for CT projections is the posterior Insula [16, 17]. Accordingly, fMRI studies have shown that affective touch elicits a velocity-specific hemodynamic response in the contralateral posterior Insula [16, 18–20], and in higher-order regions such as orbitofrontal cortices [21, 22]. The Insula is also thought to sustain the early convergence of sensory and affective signals about the body, contributing to so-called ‘interoceptive processing’ [17, 23]. According to Craig [23], interoception refers to the awareness of the physiological condition of the body, which integrates both sensations coming from within the body, (e.g. cardiac, respiratory, and digestive signals), and from the outside (e.g., temperature, itch, pain, and pleasure). In turn, the processing and integration of interoceptive information contributes to the maintenance of body awareness and to the regulation of homeostasis [24–25]. CT projections to insular brain regions characterize these afferents as an interoceptive modality, therefore conveying information about the affective and physiological state of the body, and ultimately contributing to body awareness [17, 26].

Top-down influences (e.g., the learnt affective and social meanings of interpersonal touch) can affect the hedonic value of touch leading to a high inter-individual variability in tactile pleasantness [27]. For example, the motivational style and gender of the touch receiver might influence the perceived pleasantness of interpersonal touch [28]. Differences based on individual’s social abilities, as measured by autistic traits [29] or attachment style [30], may also alter social touch perception. Moreover, recent research has shown that touch elicits lower subjective pleasantness ratings in individuals with Anorexia Nervosa (AN), a trait that endures even after an otherwise successful recovery [31, 32]. Anorexia Nervosa is a psychiatric condition characterised by profound disturbances in eating behaviours and body image [33], accompanied by impairments in social cognition and in the perception of interoceptive bodily sensations [34, 35]. These impairments might also extend to the interoceptive modality of affective touch, indicating the presence of a bodily-encoded anhedonia in interpersonal interactions [32].

Further studies have recently shown that both acute and recovered AN (RAN) patients also display a reduced responsivity to the anticipation of CT-optimal touch, both in terms of neural response localized to the ventral mid-insula, and in terms of predicted pleasantness of touch [32, 36]. Taken together, these findings suggest that in AN there is a dysregulation in the ability to correctly predict and interpret interoceptive stimuli, including CT-based pleasantness of touch.

Different lines of research have shown that body perception in Eating Disorders (EDs) is characterised by an over-investment on external physical features of the body (i.e., body image concerns) coupled with a blunted perception of information arising from within the body (i.e., interoception). Accordingly, research in clinical and non-clinical populations has shown the existence of a link between body dissatisfaction, physical appearance preoccupation and interoceptive deficits [37, 38]. Yet, this would be the case for all psychiatric conditions in which the body is a motive for concerns, including not only EDs but also, for example, Body Dysmorphic Disorder (BDD) [39, 40]. Like EDs, individuals with BDD typically exhibit a distorted image of their bodies and preoccupation with their appearance and shape [41]. Despite the fact that somatic misperceptions and low levels of interoceptive awareness have been included as part of a theoretical model of BDD [42], there are no studies which attest the putative role of CTs in such clinical conditions, a question which warrants further investigation.

The purpose of this study was threefold. First, we examined whether pleasantness of touch varies across different hairy skin body sites that are innervated by CTs. In other mammals these nerves are called C-low threshold mechanoreceptors (CLTM) and in a study with mice it was shown that CLTMs more densely innervate proximal body sites, such as the back and head, compared to distal sites [43]. In a recent study with humans where participants viewed video clips of touch to different body sites and were asked how pleasant they thought the touch was to the receiver, sites that are more proximal were also reported as being more pleasant [44]. However, the classical approach for characterising CT (actual) responses has usually limited its focus to the forearm, compared to the glabrous skin of the palm, which should be CT-free. To the best of our knowledge, only three studies have looked so far at differences in pleasantness ratings at several stroking velocities, across several skin sites [3, 4, 45], thus characterising the pleasantness/stroking velocity profile for each of these body sites. In the current study, we wanted to replicate and extend previous findings by targeting additional body sites to include the cheek, back and abdomen.

Second, we wanted to explore whether EDs and BDD traits modulate the way touch is experienced in the above-mentioned body sites. Previous psychological studies targeting AN population have mostly focussed on the somatosensory (discriminative) aspect of tactile perception only (mainly processed by large diameter, fast conducting, myelinated Aβ afferents) [46, 47]. For example, Keizer et al. [48–50] used a 2-point discrimination (2pd) task and reported that AN patients overestimate 2pd distances when presented to emotionally salient body site, such as the abdomen. Furthermore, another study by Spitoni et al. [51], which measured 2pd thresholds, found that AN patients showed relative overestimation of tactile distances on the abdomen compared to the sternum. Whilst overall these findings seem to provide evidence of an impairment of the somatosensory, discriminatory aspect of touch at emotionally salient body sites and its link to body image disturbance, no studies so far investigated whether responses to affective touch in the presence of EDs and BDD traits are modulated by specific body sites with a possible negative emotional valence. To this aim, we targeted for the first time, an emotionally salient body site with a negative valence, i.e., the abdomen, which is usually linked to worry of fatness and dissatisfaction in the general population [52–55] and in individuals experiencing EDs [56–58]. Furthermore, we targeted the face area as it represents a body site which is salient for the construction of one’s own identity and body image [59], and also because individuals suffering from BDD commonly experience concerns about their facial appearance [60, 61, 62].

Third, given the importance of body awareness in one’s sense of self, we were interested in exploring the relationship between subjective experience of pleasant touch at the different body sites with individual differences in subjective reports of interoceptive sensibility and deficits - two separate core deficits crucially associated with EDs and BDD [39, 40, 63]. To do so, we employed an affective touch manipulation task during which participants were required to provide pleasantness ratings of light, dynamic stroking touches applied to five body sites: palm, face, abdomen, back and forearm. One velocity was targeted to optimally activate CTs (3 cm/s) whereas the other two, faster (30 cm/s) and slower (0.3 cm/s) strokes fell outside the CT optimal range [3].

Furthermore, we collected ratings about participants’ touch intensity experience at the varying body sites. With these regards, recent evidence suggests subjective evaluations of touch responses, such as intensity ratings, are not velocity specific in the same way as pleasantness ratings [64, 65], and that recovered anorexics display lower insular cortex responses during touch anticipation associated with higher perceived touch intensity ratings [36]. Therefore, we collected participants’ ratings of touch intensity, to account for potential individual differences in the two aspects of touch (discriminative vs. affective).

Overall, we expected that differences in touch velocities and application sites would inform the degree to which responses to predominantly discriminatory (touch to the palm and non-CT optimal touch) compared to affective/CT-activating (e.g., CT targeted 3 cm/s touch to the cheek, abdomen, back and forearm) tactile stimuli are associated with dysmorphic appearance concerns and body awareness. Based on previous research investigating the relationship between touch pleasantness and stroking velocity at the forearm [3], we predicted that participants would prefer the CT-optimal (3cm/s) stroking velocity, with reduced pleasantness for non-CT optimal (0.3 cm/s) slower stroking velocity and the non-CT optimal (30 cm/s) faster stroking velocity. Crucially, based on previous findings from clinical populations [31, 66], we also expected to observe a significant relationship between dysmorphic appearance concerns and pleasant touch awareness, particularly at emotionally salient body sites (for e.g., abdomen and face, as compared to the palm). Finally, we hypothesised that body awareness and interoceptive deficits would be related to pleasant touch awareness perception, so that individuals’ (hyper)sensitivity to internal bodily sensations and interoceptive deficits will be predictive of participants’ preference of CT-optimal (3cm/s) touch. At the same time, the sensory, touch intensity, aspect of touch was expected to differ across the different body sites and to be associated to individual differences in dysmorphic appearance concerns and body awareness.

## Methods

### Participants

The sample size was based on a preliminary calculation using the freely available G*Power software (G*Power 3.1.9) [67], which indicated a minimum sample of 24 participants as adequate for a design with 95% power to detect a moderate effect size (*f* = 0.25), using a repeated-measures ANOVA with alpha at .05 (two tailed). To cope with potential drop out and outlier case exclusion, a total of 30 women (mean age = 25.87±1.17yrs; mean BMI = 27.22±1.06kg/m^2^) were recruited.

Participants were recruited internally through the Liverpool John Moores University (LJMU) research participation system for undergraduate Psychology students in exchange for course credits or externally through poster advertisements situated in public locations, social media and through individuals known to the researchers. In line with previous literature analysing body perception and eating behaviours, the current study employed a population of only female participants [53, 68, 69]. Indeed, literature on the prevalence and the phenomenology of EDs in male populations is still limited [70]. All participants, except one, were right-handed as assessed by the Edinburgh Handedness Inventory [71]. All reported normal or corrected to normal vision and they were in good health, free of psychotropic or vasoactive medication, with no current or history of any psychiatric or neurological disease, no skin condition (e.g. psoriasis, eczema, etc.) and not pregnant. Participants provided written informed consent prior to testing and were debriefed at the end of the experiment. All procedures were approved by the LJMU research ethics committee, in agreement with the ethical standards of the 1964 Declaration of Helsinki.

### Self-report questionnaires

#### Eating Disorder Inventory-3

The Eating Disorder Inventory-3 (EDI-3) [72] comprises 91 items assessing ED core symptoms (i.e., Drive for Thinness, Bulimia and Body Dissatisfaction), as well as other psychological constructs and personality traits that have been associated with disordered eating behaviours (Low Self-esteem, Personal Alienation, Interpersonal Insecurity, Interpersonal Alienation, Interoceptive Deficits, Emotional Dysregulation, Perfectionism, Ascetism and Maturity Fear). Examples of items of this scale include items such as “I think my hips are too big”, “I am preoccupied with the desire to be thinner” “I feel alone in the world” and “I stuff myself with food”. Participants are instructed to rate each item on a 6-point Likert scale ranging from ‘never’ to ‘always’. The EDI-3 allows the calculation of 12 scales and 6 composites describing different aspects of the ED symptomatology.

In the current study, we were specifically interested in assessing whether participants’ overall level of ED symptoms and interoceptive abilities could influence their experience of pleasantness and intensity related to touch. To this aim, we focussed our analyses on the ED Risk Composite (EDRC) and the Interoceptive Deficits subscale. The EDRC score represents an index of the risk to develop an ED and is calculated by summing the scores of the three symptom-specific subscales of the inventory: Drive for Thinness, Bulimia and Body Dissatisfaction. The Interoceptive Deficits subscale is an index of participants’ self-reported ability to correctly recognize and respond to inner bodily states. The EDI-3 was validated in clinical and non-clinical samples, and has shown good internal consistency (α = between .75 and .92 for each subscale), and excellent sensitivity and specificity [73].

### Dysmorphic Concern Questionnaire

The Dysmorphic Concern Questionnaire (DCQ) [74] comprises 7 items investigating participants’ concern about their physical appearance. The scale has been validated as a brief, sensitive, and specific screening measure of trait-level BDD symptoms [75]. Items focus on the belief of being misshapen or malformed despite others’ opinion; belief in bodily malfunction (e.g. malodour); consultation with cosmetic specialists; spending excessive time worrying about appearance; and spending a lot of time covering up perceived defects in appearance. Participants were asked to rate each item on a Likert scale from a minimum of 0 (“not at all”) to a maximum of 4 (“much more than most people”). Total scores range from 0 to 28 with a critical value of 9 indicating clinical concern [75]. The scale was administered to investigate whether individual differences in physical appearance concerns (higher trait BDD symptoms) could explain participants’ responses to CT-optimal touch at the different body sites. The DCQ has been shown good internal consistency with α= .80 [76].

### Body Perception Questionnaire-Short Form

The Body Perception Questionnaire-Short Form (BPQ-SF) [77] is a 46-item self-report questionnaire for the assessment of participants’ awareness about different bodily states associated with changes in the activity of the autonomic nervous system, such as “muscle tension”, “goose bumps”, “stomach and gut pains”, breathing and heart-beat rates. Items are rated on a 5-point Likert scale ranging from 1 (“Never”) to 5 (“Always”), and total scores ranging between 12 and 60. In the current study, we focussed on the Body Awareness subscale of the BPQ (26 items) which reflects participants’ sensitivity to inner bodily sensations, with higher values reflecting a *hypersensitivity* and lower values a *hyposensitivity* to bodily sensations. Together with the Interoceptive Deficit subscale of the EDI-3, the BPQ was administered to investigate whether individual differences in body awareness could explain differences in participants’ responses to CT-optimal touch at different body sites. The BPQ has been validated in different samples and has been shown to have good internal consistency (α = between .88 and .97), and an excellent test-retest reliability [78].

### Body Mass Index

Each participant’s body mass index (BMI) was physically calculated on the day of the experimental session, from their weight and height obtained from a calibrated digital scale (OMRON BF511) and a stadiometer.

### General Procedure

Participants were lying semi-horizontally in a comfortable, reclining chair whilst receiving manual brush strokes. Although during each trial participants kept their eyes open, the position on the chair prevented participants from seeing the stroking procedure.

During the experiment, participants received manual brush strokes to the dorsal surface of their forearm, cheek, palm, upper part of the back and stomach site of their right body using a soft cosmetic brush (No7 cosmetic brush, Boots UK). To avoid any social confound, stroking touch was delivered by the same female experimenter to all participants (SSh).

During each trial, strokes were delivered for 6 secs on a 9 cm surface at one of three set velocities: 0.3 cm/s, 3 cm/s, and 30 cm/s. The experimenter was trained to deliver stroking touch with a constant pressure of 220 mN, which was calibrated using a high precision digital scale. The stroking was delivered in a proximal to distal direction.

Overall, the stroking task consisted of 5 blocks: one block for each body site. Each block consisted of 9 trials, with 3 trials for each velocity. Across the 5 blocks, participants experienced a total of 15 CT-optimal trials (3 cm/s, 2 strokes per trial), 15 non-CT optimal fast touch trials (30 cm/s, 20 strokes per trial) and 15 non-CT optimal slow touch trials (0.3 cm/s, 1/5 of a stroke per trial corresponding to 1.8 cm). Participants were randomly assigned to receive the experimental blocks in one of 5 pseudo-randomised orders, which ensured no two consecutive trials were the same.

The running orders were created in PsychoPy [79]. Trial velocity was randomised to minimize habituation [80]. Trials were delivered 30s apart, as CT fibers are easily fatigued [9].

A visual metronome, programmed in PsychoPy, was presented on a computer screen behind the participant [81, 82] and guided the researcher in delivering the brush strokes at one of the three velocities.

After each trial participants rated, on a paper, the pleasantness and intensity of the stimulation on 15 cm horizontal Visual Analogue Scales (VAS) ranging respectively from “unpleasant” to “pleasant”, and from “least intense” to “most intense” (see Fig 1).

**Fig 1:**
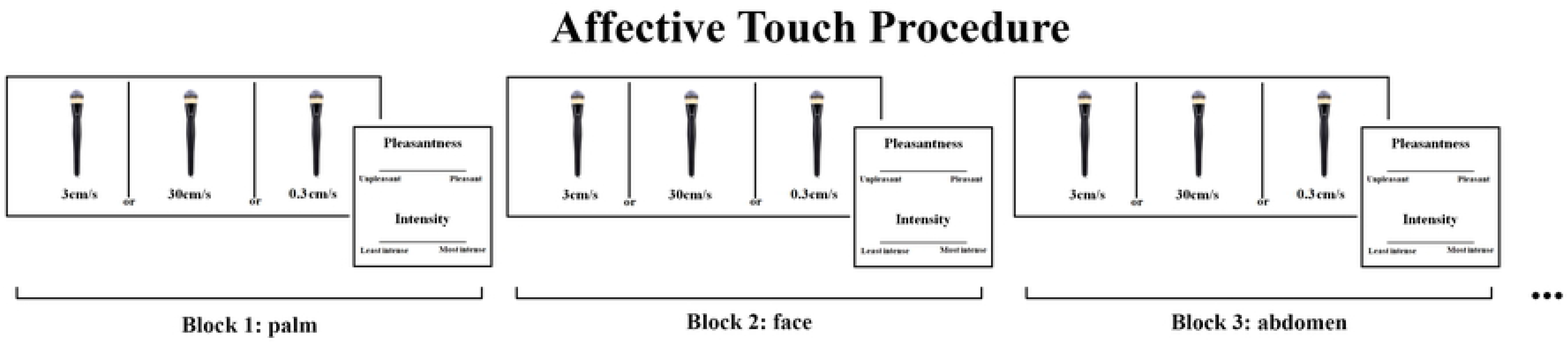
Schematic depiction of the study procedure. Participants were lying semi-horizontally in a comfortable, reclining chair whilst receiving manual brush strokes. In randomised blocks (each one corresponding to one body site), participants were asked to rate the pleasantness and intensity of the touch delivered by a female experimenter to the palm, face, abdomen, back and forearm at 3 cm/s (CT-optimal stroking) and 0.3 cm/s and 30 cm/s (non-CT optimal stroking) velocities.

At the end of the experiment, participants were weighted on the digital scale and completed the self-report questionnaires.

### Data Handling

All statistical analyses were performed using STATISTICA 8.0 (StatSoftInc, Tulsa, Oklahoma). In our main analysis, mean pleasantness and intensity ratings were entered into two separate two-way 5×3 repeated-measures Analysis of Variance (ANOVA) with body sites (palm, face, abdomen, back and forearm) and stroking velocity (0.3 cm/s, 3 cm/s, 30 cm/s) as within-subjects factors. All data are reported as Mean (M) and Standard Error of the Mean (S.E.M.). A significance threshold of *p* < 0.05 was set for all effects and effect sizes were estimated using the partial eta square measure (*ηp*^2^). Duncan post-hoc tests were performed to follow-up significant interactions. Furthermore, following previous studies [6, 83], we computed two main affective touch indices, as follows: 1) overall touch intensity which is calculated as the mean of all touch intensity ratings and 2) pleasant touch awareness (PTA) which is calculated as the difference of pleasantness ratings between CT-optimal (3 cm/s) and non-CT optimal (30 cm/s) fast stroking, weighted by the mean of all touch pleasantness ratings (overall touch pleasantness), [PTA=(pleasantness ratings at 3 cm/s-pleasantness ratings at 30 cm/s)/overall touch pleasantness]. A positive PTA index indicates that participants prefer stroking at CT-optimal 3 cm/s velocity to stroking at non-CT optimal (30 cm/s) fast velocity. To explore the associations of these two indices with individual scores obtained at the self-report scales: EDRC (EDI-3), DCQ, Interoceptive Deficits (EDI-3) and Body awareness (BPQ), a series of multiple linear regressions were fit to predict the two measures of touch experience at each of the five body sites.

## Results

### Demographics and self-report scales

Participants’ demographics and self-report questionnaire scores are reported in Table 1. The mean EDRC score was 18.53 (±1.41, range: 1-29) which is deemed to be at low risk for the development of an ED [73]. The mean DCQ score was 7.67 (±0.80, range: 1-16) which is below the critical level of high dysmorphic concerns, with scores above 9 indicating clinical levels of body image concerns [74].

**Table 1.** Participants’ means and Standard Error of Means (S.E.M. in brackets) of demographic variables and self-report questionnaire scores.

|  | Mean (s.e.m.) | Minimum | Maximum |
| --- | --- | --- | --- |
| Age (yrs) | 25.87 (1.17) | 21 | 40 |
| BMI (kg/cm<sup>2</sup>) | 27.22 (1.06) | 18 | 40.7 |
| **EDI-3** |  |  |  |
| Drive for Thinness | 8.13 (0.78) | 0 | 16 |
| Body Dissatisfaction | 7.33 (0.64) | 0 | 12 |
| Bulimia | 3.07 (0.45) | 0 | 8 |
| Low self esteem | 1 (0.21) | 0 | 4 |
| Personal Alienation | 5.83 (0.76) | 0 | 15 |
| Interpersonal Insecurity | 7.3 (0.92) | 0 | 17 |
| Interpersonal Alienation | 6.7 (0.91) | 0 | 20 |
| Interoceptive Deficit | 7.57 (1.10) | 0 | 24 |
| Emotional Dysregulation | 8.03 (1.14) | 0 | 24 |
| Perfectionism | 10.03 (1.12) | 0 | 23 |
| Ascetism | 7 (1.01) | 0 | 21 |
| Maturity Fear | 9.67 (1.16) | 1 | 28 |
| EDCR | 18.53 (1.41) | 1 | 29 |
| DCQ | 7.67 (0.80) | 1 | 16 |
| Body Awareness (BPQ) | 63.17 (4.13) | 29 | 112 |

**Notes:** *BMI*, Body Mass Index; *EDI-3*, Eating Disorder Inventory-3; *EDCR*, Eating Disorder Risk Composite; *DCQ*, Dysmorphic Concern Questionnaire; *BPQ*, Body Perception Questionnaire.

### Main Analysis

#### Pleasantness Ratings

The 2-way ANOVA of pleasantness ratings revealed a significant main effect of body sites [*F*(4,116) = 4.475, *p* = 0.002, ηp2 = 0.134] with touch to the abdomen rated significantly lower (all *ps* < 0.047) than any other body site. No significant differences were observed amongst all other body sites (all *ps*> 0.311). There was also a significant main effect of stroking velocity [*F*(2,58) = 18.706, *p* < 0.001, *ηp2* = 0.392]. Whilst the touch applied at CT-optimal (3 cm/s) stroking velocity was rated significantly more pleasant than the touch applied at the other two non-CT optimal velocities (all *ps* < 0.001), no difference was observed between the two non-CT optimal (0.3 cm/s vs. 30 cm/s) stroking velocities (*p*=0.593). Importantly, the 2-way interaction of body sites and stroking velocity was also significant [*F*(8,232) = 2.594, *p* = 0.010, *ηp^2^* = 0.082, see Fig 2]. Post-hoc comparisons showed that whilst there was no difference amongst the palm, face, back and forearm body sites for the preferred CT-optimal (3 cm/s) stroking velocity (all *ps* > 0.314), touch delivered at CT-optimal (3cm/s) velocity to the abdomen was rated significantly less pleasant compared to all body sites (all *ps* < 0.001). Moreover, whilst participants rated the touch delivered at the non-CT optimal (30 cm/s) velocity as similarly less pleasant than the non-CT optimal (0.3 cm/s) touch at all body sites (all *ps* > 0.697), this was not the case for the abdomen. At this body site, the non-CT optimal (30 cm/s) touch was rated as significantly less pleasant than the non-CT optimal (0.3 cm/s) touch (*p* = 0.029). However, this effect was not specific for the abdomen, since no significant difference in pleasantness was observed between the abdomen and the face, when the stroking was delivered at non-CT optimal (30cm/s) velocity (*p*=0.407). Furthermore, no difference for the non-CT optimal (30cm/s) touch was observed when comparing this velocity amongst the palm, back and forearm (all *ps*>0.111). However, participants rated the non-CT optimal (30cm/s) touch as less significantly pleasant when delivered to the face and the abdomen (all *ps*<0.045). Finally, no significant differences in pleasantness were observed when comparing non-CT Optimal (0.3 cm/s) touch delivered at all body sites (all *ps*>0.394).

**Fig 2:**
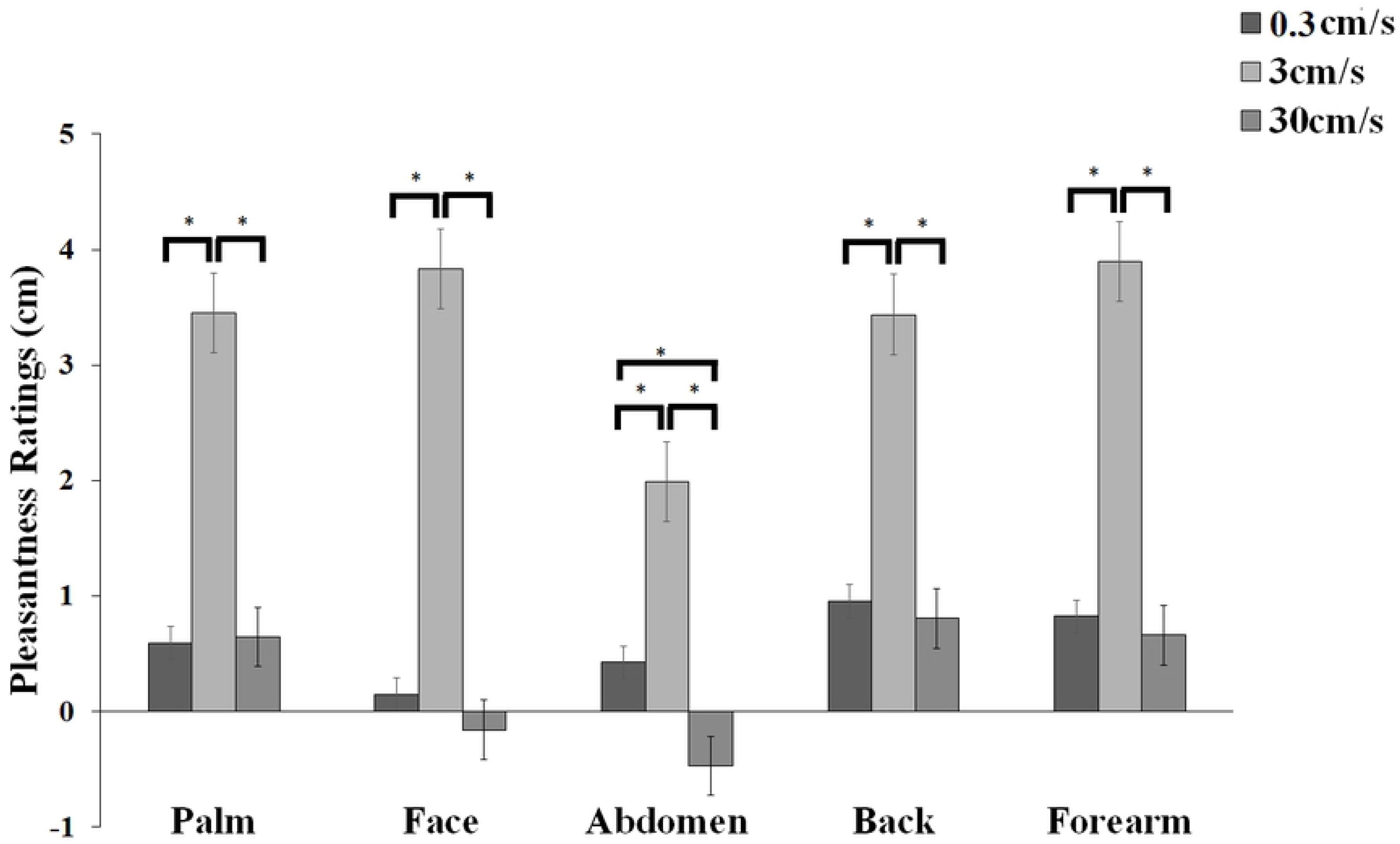
Mean Pleasantness Ratings (+/- s.e.m) for touch delivered at each of five body sites (Palm, Face (cheek), Abdomen, Back, Forearm) at 3 cm/s (CT-optimal stroking) and 0.3 cm/s and 30 cm/s (non-CT optimal stroking) velocities.

Additionally, separate regression analyses for each body site were carried out to examine whether the relationship between perceived pleasantness and stroking velocities was better described by linear or quadratic models. Results consistently showed at every body site, profiles were all best fit by negative quadratic models (all *ps* <0.001), rather than linear models (all *ps* > 0.20), giving the characteristic “inverted-U” shaped curves, as found in previous studies investigating pleasantness of different velocity stroking stimuli [3, 45].

### Intensity Ratings

The 2-way ANOVA of intensity ratings revealed a significant main effect of body sites [*F*(4,116) = 6.203, *p* < 0.001, ηp2 = 0.176], with touch to the back rated significantly less intense than any other body site (all *ps* < 0.044). No statistical difference in intensity ratings was identified between the other body sites (all *ps*>0.080). Furthermore, and as expected, the main effect of stroking velocity was significant [*F*(2,58) = 35.705,*p* < 0.001, ηp^2^ = 0.552] with non-CT optimal (30 cm/s) touch being rated as significantly more intense than the CT-Optimal (3 cm/s) and non-CT optimal (0.3 cm/s) touch (all *ps* < 0.001). No statistical difference was observed between the intensity ratings for the non-CT optimal (0.3 cm/s) touch and the CT-optimal (3cm/s) touch (*p*= 0.874). The 2-way interaction was not significant [*F*(8,232) = 0.761, *p* = 0.637, *ηp^2^* = 0.026, see Fig 3].

**Fig 3:**
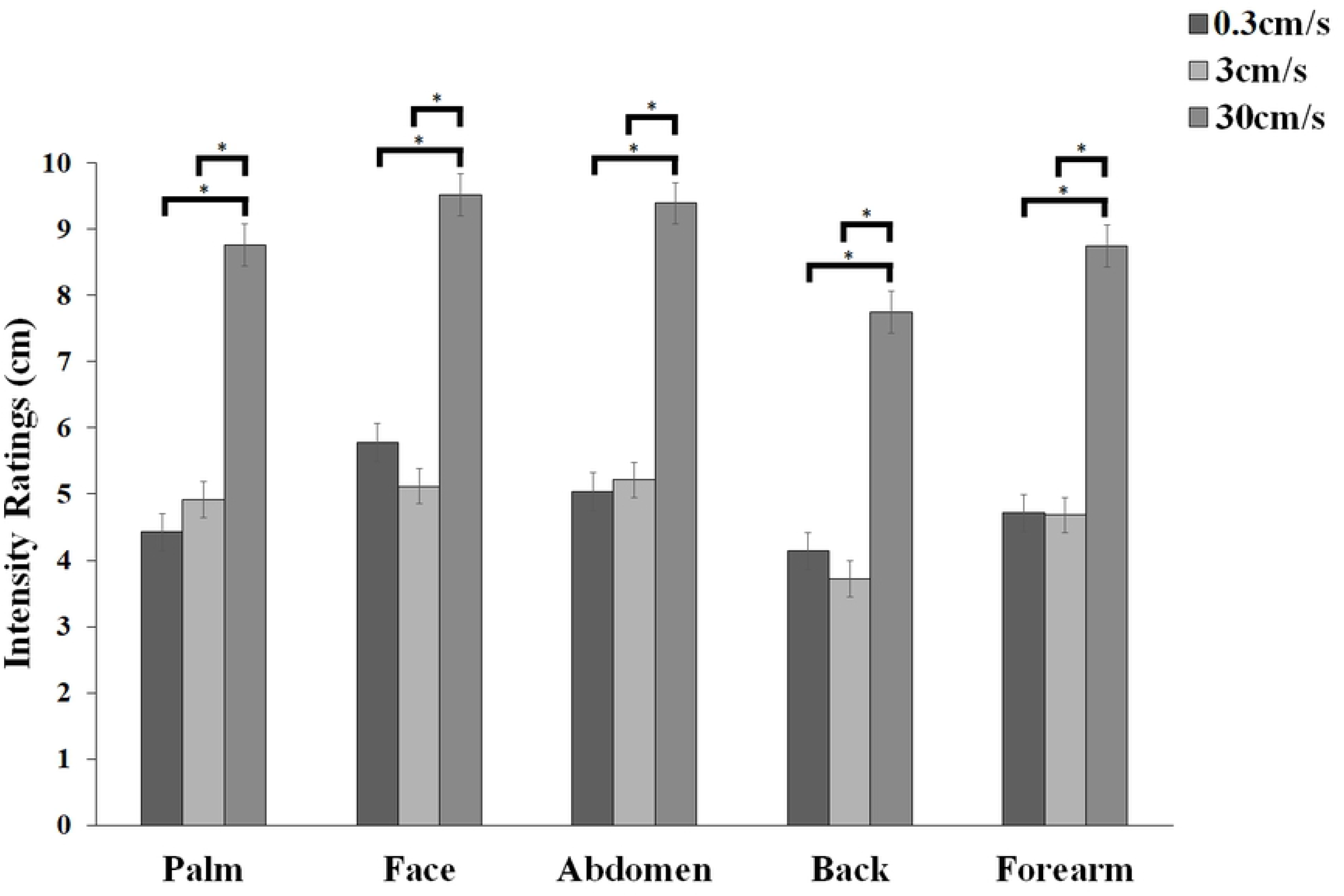
Mean Intensity Ratings (+/- s.e.m) for touch delivered at each of five body sites (Palm, Face (cheek), Abdomen, Back, Forearm) at 3 cm/s (CT-optimal stroking) and 0.3 cm/s and 30 cm/s (non-CT optimal stroking) velocities.

### Multiple regression analyses: affective touch indices and eating and dysmorphic concerns symptomatology

By means of a series of exploratory (due to the relatively small sample size for such analyses) multiple regression analyses, we assessed whether symptoms of eating and dysmorphic concerns, interoceptive deficits and body awareness could predict participants’ PTA and overall touch intensity at all body sites. To this aim, we ran separate multiple regressions to explore whether EDRC (EDI-3), DCQ, interoceptive deficits (EDI-3) and Body Awareness (BPQ) were significant predictors of PTA and overall touch intensity, at each body site.

The linear multiple regression analysis calculated to predict PTA for the back from symptoms of eating and dysmorphic concerns, interoceptive deficits and Body Awareness scores was significant [*F*(4,25) = 2.959, *p* = 0.039, *R*^2^=0.321], with Body Awareness emerging as the only significant predictor (see Table 2). Furthermore, we found a significant regression equation for the forearm with DCQ and Interoceptive deficit scores being significant predictors [*F*(4,25) = 3.770, *p* = 0.016, *R^2^* = 0.376, see Table 2]. We found non-significant regression equations for the palm [*F*(4,25) = 0.787, *p* = 0.544, *R*^2^=0.112], face [*F*(4,25) = 1.204, *p* = 0.334, *R*^2^=0.161], and abdomen [*F*(4,25) = 0.407, *p* = 0.801, *R*^2^=0.061]. Full results are reported in Table 2.

**Table 2:** Unstandardized coefficients from the linear multiple regression model of eating and dysmorphic concerns and body awareness predictors of the pleasantness touch awareness (PTA) index at each body site (palm, face, abdomen, back and forearm).

| Pleasantness Touch Awareness |  |  |  |  |  |
| --- | --- | --- | --- | --- | --- |
| Palm |  |  |  |  |  |
| | B | SE | $\beta$ | *t*-value | *p*-level |
| **EDRC (EDI-3)** | 0.14 | 0.21 | 0.05 | 0.65 | 0.52 |
| **DCQ** | 0.09 | 0.26 | 0.05 | 0.33 | 0.75 |
| **Interoceptive Deficit**<br>**(EDI-3)** | 0.09 | 0.23 | 0.04 | 0.39 | 0.70 |
| **Body Awareness (BPQ)** | -0.31 | 0.21 | -0.04 | -1.47 | 0.15 |
| Face |  |  |  |  |  |
| | B | SE | $\beta$ | *t*-value | *p*-level |
| **EDRC (EDI-3)** | -0.20 | 0.21 | -0.13 | -0.95 | 0.35 |
| **DCQ** | -0.27 | 0.25 | -0.31 | -1.04 | 0.31 |
| **Interoceptive Deficit**<br>**(EDI-3)** | 0.38 | 0.23 | 0.33 | 1.69 | 0.10 |
| **Body Awareness (BPQ)** | -0.14 | 0.21 | -0.03 | -0.67 | 0.51 |
| Abdomen |  |  |  |  |  |
| | B | SE | $\beta$ | *t*-value | *p*-level |
| **EDRC (EDI-3)** | -0.13 | 0.22 | -0.09 | -0.57 | 0.57 |
| **DCQ** | -0.22 | 0.27 | -0.30 | -0.83 | 0.41 |
| **Interoceptive Deficit**<br>**(EDI-3)** | 0.19 | 0.24 | 0.18 | 0.78 | 0.44 |
| **Body Awareness (BPQ)** | -0.01 | 0.22 | -0.00 | -0.04 | 0.97 |
| **Back** |  |  |  |  |  |
| | B | SE | $\beta$ | *t*-value | *p*-level |
| **EDRC (EDI-3)** | -0.05 | 0.19 | -0.01 | -0.24 | 0.81 |
| **DCQ** | -0.17 | 0.23 | -0.10 | -0.76 | 0.45 |
| **Interoceptive Deficit**<br>**(EDI-3)** | 0.02 | 0.20 | 0.01 | 0.09 | 0.93 |
| **Body Awareness (BPQ)** | **-0.46** | **0.19** | **-0.05** | **-2.47** | **0.02\*** |
| **Forearm** |  |  |  |  |  |
| | B | SE | $\beta$ | *t*-value | *p*-level |
| **EDRC (EDI-3)** | 0.06 | 0.18 | 0.02 | 0.33 | 0.75 |
| **DCQ** | **-0.62** | **0.22** | **-0.38** | **-2.82** | **0.01\*** |
| **Interoceptive Deficit**<br>**(EDI-3)** | **0.51** | **0.20** | **0.23** | **2.61** | **0.01\*** |
| **Body Awareness (BPQ)** | -0.20 | 0.18 | -0.02 | -1.10 | 0.28 |

**Notes:** EDRC, Eating Disorder Risk Composite; EDI-3, Eating Disorder Inventory-3; BPQ, Body Perception Questionnaire. \* indicates $p < 0.05$

The linear multiple regression analyses calculated to predict overall touch intensity from symptoms of eating and dysmorphic concerns, interoceptive deficits and body awareness scores were non-significant for all body sites [Palm: [*F*(4,25) = 0.652, *p* = 0.631, *R*^2^=0.094], Face [*F*(4,25) = 0.306, *p* = 0.871, *R*^2^=0.047], Abdomen [*F*(4,25) = 0.315, *p* = 0.865, *R*^2^=0.048], Back [*F*(4,25) = 0.755, *p* = 0.564, *R*^2^=0.108] and Forearm [*F*(4,25) = 0.953, *p* = 0.450, *R*^2^=0.132]. Full results are reported in Table 3.

**Table 3:** Unstandardized coefficients from the linear multiple regression model of eating and dysmorphic concerns and body awareness predictors of the overall touch intensity index at each body site (palm, face, abdomen, back and forearm).

| Overall Touch Intensity |  |  |  |  |  |
| --- | --- | --- | --- | --- | --- |
| Palm |  |  |  |  |  |
| | B | SE | $\beta$ | *t*-value | *p*-level |
| EDRC (EDI-3) | 0.19 | 0.22 | 0.06 | 0.86 | 0.40 |
| DCQ | 0.14 | 0.26 | 0.08 | 0.52 | 0.61 |
| Interoceptive Deficit<br>(EDI-3) | 0.05 | 0.24 | 0.02 | 0.21 | 0.83 |
| Body Awareness (BPQ) | 0.00 | 0.22 | 0.00 | 0.01 | 0.99 |
| Face |  |  |  |  |  |
| | B | SE | $\beta$ | *t*-value | *p*-level |
| EDRC (EDI-3) | 0.11 | 0.22 | 0.04 | 0.51 | 0.62 |
| DCQ | 0.18 | 0.27 | 0.11 | 0.65 | 0.52 |
| Interoceptive Deficit<br>(EDI-3) | -0.03 | 0.24 | -0.01 | -0.11 | 0.91 |
| Body Awareness (BPQ) | -0.09 | 0.22 | -0.01 | -0.40 | 0.69 |

| **Abdomen** |  |  |  |  |  |
| --- | --- | --- | --- | --- | --- |
| | B | SE | $\beta$ | *t*-value | *p*-level |
| **EDRC (EDI-3)** | -0.04 | 0.22 | -0.02 | -0.20 | 0.84 |
| **DCQ** | -0.08 | 0.27 | -0.05 | -0.28 | 0.78 |
| **Interoceptive Deficit**<br>**(EDI-3)** | 0.26 | 0.24 | 0.13 | 1.06 | 0.30 |
| **Body Awareness (BPQ)** | -0.08 | 0.22 | -0.01 | -0.36 | 0.72 |
| **Back** |  |  |  |  |  |
| | B | SE | $\beta$ | *t*-value | *p*-level |
| **EDRC (EDI-3)** | -0.15 | 0.22 | -0.04 | -0.68 | 0.50 |
| **DCQ** | 0.25 | 0.26 | 0.13 | 0.95 | 0.35 |
| **Interoceptive Deficit**<br>**(EDI-3)** | 0.21 | 0.23 | 0.08 | 0.89 | 0.38 |
| **Body Awareness (BPQ)** | -0.14 | 0.21 | -0.01 | -0.63 | 0.53 |
| **Forearm** |  |  |  |  |  |
| | B | SE | $\beta$ | *t*-value | *p*-level |
| **EDRC (EDI-3)** | -0.07 | 0.21 | -0.02 | -0.31 | 0.76 |
| **DCQ** | 0.28 | 0.26 | 0.16 | 1.07 | 0.30 |
| **Interoceptive Deficit**<br>**(EDI-3)** | 0.20 | 0.23 | 0.09 | 0.88 | 0.38 |
| **Body Awareness (BPQ)** | -0.13 | 0.21 | -0.01 | -0.63 | 0.53 |

**Notes:** EDRC, Eating Disorder Risk Composite; EDI-3, Eating Disorder Inventory-3; BPQ, Body Perception Questionnaire.

## Discussion

This study aimed to: a) investigate whether pleasantness and intensity ratings of touch vary across different body sites, b) explore associations between EDs and BDD traits with gentle touch applied at several body sites and in particular at emotionally salient body sites, i.e., abdomen and face; c) explore the relationship between body awareness and interoceptive deficits with tactile experience at these varying body sites.

The results show that, as expected, perception of touch varied across skin sites according to both tactile pleasantness and intensity. Tactile pleasantness, measured by means of hedonic ratings to three velocities of stroking, one CT-optimal (3 cm/s) and two non-CT optimal (0.3 cm/s and 30 cm/s), was similar in profile across the body sites investigated. However, we found that there was a subtle difference in the ratings at the abdomen, so that participants rated this body area the least pleasant when touch was delivered at CT-optimal (3cm/s) velocity. However, this reduction in pleasantness for CT-optimal touch to the abdomen was not supported by a parallel increase in the intensity sensation experienced by participants at the same body site, given that overall, touch to the back but not to the abdomen was felt as the least intense. Furthermore, and contrary to our prediction, we did not find evidence to support the idea of an association between preference for CT-optimal (3 cm/s) touch when applied to the abdomen and individual differences in any of the affective body image components. Interestingly however, exploratory multiple regression analyses showed that perceived tactile preference for CT-optimal (3 cm/s) touch delivered to the back, was predicted by individual differences in body awareness (BPQ). Similarly, dysmorphic concerns and interoceptive deficits emerged as significant predictors for perceived pleasantness of CT-optimal (3 cm/s) touch to the forearm. We will now proceed to discuss specific findings in turn.

As expected, our main analysis of pleasantness ratings showed that participants expressed higher ratings for the CT-optimal (3 cm/s) velocity, compared to the slower (0.3 cm/s) or faster CT-non optimal (30 cm/s) stroking, at all body sites. This finding was corroborated by the regression analysis, which showed that a negative quadratic fit best described the stroking velocity-pleasantness profiles, giving the characteristic “inverted-U” shaped curves, at all body sites.

Essick et al. [45] found a similar preference for stroking at 5 cm/s on the cheek, compared to slower (0.5 cm/s) and faster (50 cm/s) strokes, and Essick et al. [3] showed that pleasantness was higher for stroking at 5 cm/s than at 20 cm/s at the forehead, finger (glabrous skin), thigh and calf. In partial agreement with current findings, we also found that (except for the abdomen area), pleasantness ratings at the glabrous, CT-free palm were not different from other body sites, where CTs are found. This was also supported by the result that a quadratic function best explained the data for relative touch pleasantness when applied to the glabrous palm.

Although numerous studies compare the CT-free palm to other hairy skin sites where CTs are present (see a recent review [11]), it is clear that pleasantness in touch can be experienced at glabrous skin sites, despite its reported lack of CTs [83, 84]. With this regard, and in accordance to our results, Löken et al. [3, 83] found that an ‘inverted-U’ velocity profile was present in pleasantness ratings for the palm, whist the overall mean was lower compared to stroking on the arm. One plausible explanation for these findings is that not only peripheral input, but also top-down mechanisms might have modulated the perception of pleasant touch at the glabrous palm [85, 86]. For instance, several studies have shown that touch stimulation of both hairy and glabrous skin similarly activates the orbitofrontal cortex [20–22], a brain area which is important for linking affective experiences to its rewarding value [86]. Therefore, as suggested by McGlone et al. [87], it may be plausible that the sensation of pleasantness at the palm is likely produced by the combination of unmyelinated and myelinated mechanoreceptive input, where higher-order cognitive evaluation of this type of touch is paramount in the judgment of its hedonic value.

In the present work and as predicted, we found a reduction in subjective reports of pleasantness when CT-optimal (3 cm/s) touch was delivered to the abdomen. Particularly, the abdomen was felt as less pleasant compared to all other body sites, and was rated significantly less pleasant when touch was applied at the faster velocity (30 cm/s) compared to CT-optimal (3 cm/s) and slower (0.3 cm/s) touch. The effect was specific for the abdomen, given the significant differences observed when comparing the mean pleasantness ratings for CT-optimal touch to all other body sites. However, when we explored the relationship between pleasantness touch awareness (PTA) for the abdomen and individual differences in EDs and BDD traits, as well as body awareness/deficits no evidence for a linear association was found with such variables. This suggests that the reduction in perceived pleasantness when CT-optimal (3 cm/s) touch is delivered to this body area is not explained by participants’ EDs or BDD traits, nor by their hypersensitivity to bodily signals or interoceptive deficits.

We know from previous literature that the abdomen area has been identified as the body site women generally perceive as least attractive/satisfied with [52, 53, 88, 89], and that this body site is often subject to tactile overestimation in individuals suffering from EDs [48, 51]. However, one alternative explanation to account for this overall reduction in perceived pleasantness at the abdomen may reside in the fact that for women the meaning of a touch is primarily influenced by how well the touch receiver knows the other, with touch from a same-sex person rated as unpleasant [90]. With this regard, a recent study from Suvilehto and colleagues [91] reported that across a broad range of European countries, whilst emotionally closer individuals in inner layers of the social network (e.g., family members) were allowed to touch wider body areas, touching by strangers was primarily limited to the hands and upper torso. However, the same study also reported that females, rather than males, evaluate touch as more pleasant and that they allow themselves to be touched on a larger body area than males (see also [92]). Furthermore, female same-sex touch was allowed without discomfort on most of the body’s surface [91]. Although, our sample of women do not rate CT-optimal (3 cm/s) touch applied to the abdomen as extremely unpleasant and evidence reported above seems to speak against this, at present we cannot rule out if this reduction in perceived pleasantness might be the result of a subjective feeling of intrusiveness of touch, as applied by the same-sex, stranger experimenter.

An additional but not mutually exclusive explanation of the lack of association between reduced pleasantness at the abdomen and EDs/BDD traits may reside in the fact that in our sample there was essentially no evidence for particularly high-risk ED psychopathology, as shown by a relatively low spread and mean of EDRC (EDI-3) scores. Specifically, our sample of women scored an average of 18.53 on the EDRC, which is deemed to be ‘at low risk’ for the development of an ED [73]. We acknowledge this is a limitation to our study and that future research is necessary to better address whether clinical populations suffering from EDs and BDD may show reduced pleasantness of touch when delivered to emotionally salient body areas, such as the abdomen.

Interestingly, a further unexpected finding in our study may begin to elucidate a link between interoceptive affective touch and BDD traits. Indeed, a multiple regression analyses showed that subjective ratings of pleasantness (as indexed by PTA) when delivered to the forearm were predicted by individual differences in dysmorphic appearance concerns, so that the higher women’s dysmorphic concerns the lower their preference for CT-optimal (3 cm/s) touch. Although, this result should be interpreted with caution due to its exploratory nature, this is the first study, to the best of our knowledge, to provide evidence of a relationship between a preference for CT-optimal (3 cm/s) touch and BDD symptomatology. To date, no studies so far have attempted to identify a link between BDD subthreshold symptomatology and pleasant touch. However, other studies have proposed that individuals with BDD may display lower levels of interoceptive awareness compared to individuals without this disorder [39]. Furthermore, research evidence suggests that a disconnection with the internal body may contribute to the misperception of and pre-occupation with features of the external body [40, 93]. Accordingly, our study provides first evidence that lower levels of dysmorphic appearance concerns may be linked to a preferential response to affective touch, similar to that observed by investigations into other disorders like AN, which commonly share symptoms of body image disturbance [31, 36].

Interestingly, we also found evidence that PTA (for the back and forearm) was predicted by differences in body awareness and interoceptive deficits, respectively. These results resonate with previous findings from Cabrera et al. [78], who found that BPQ scores were elevated in participants with a self-reported psychiatric diagnosis, as well as with previous clinical observations showing altered interoceptive functions across a range of psychiatric diagnoses, including Autism Spectrum Disorder, Post-Traumatic Stress Disorder [94], Anxiety [95, 96] and EDs [97].

With these regards, Pollatos et al. [34] proposed that individuals with higher levels of interoceptive awareness are likely to display heightened reactions to emotional stimuli, as they are more in tune with their bodily feedback. Furthermore, several authors have suggested that this increased awareness of somatic bodily sensations, and their subsequent appraisal as a threat to the self, contribute to the development and maintenance of anxiety disorders [95, 98]. Accordingly, we suggest that hyper-sensitivity to somatic bodily sensations (as measured by the BPQ), coupled with a greater sensitivity to CT-optimal touch may contribute all together to the aetiology and maintenance of those psychiatric conditions characterised by body image disturbance, a hypothesis that requires further exploration.

Contrary to our expectations, however, we did not find evidence of a link between overall touch intensity and self-report measures of EDs, BDD, BPQ and interoceptive deficits. This finding speaks against that reported by Bischoff-Grethe et al. [36] according to which ill and recovered anorexics reported gentle touch as more intense, thus supporting a generalized finding of elevated intensity perception to somatosensory stimuli in both ill and weight-restored AN [99]. Future research should aim to clarify whether a reduced responsivity to affective interoceptive stimuli coupled with an elevated intensity perception to somatosensory stimuli is linked to EDs and BDD traits. This would lend support to the idea that blunted interoception is linked to the (sensory) misperception of the body and its features, with both reported as being observed in both of these clinical conditions.

In conclusion this study found that overall, subjective experience of pleasant touch and intensity sensation varies across body sites, as we have already shown, but importantly, our findings begin to elucidate the unique associations between an increased sensitivity to affective interoceptive stimuli with body awareness and dysmorphic appearance concerns, thus proving the importance of extending research on affective touch to other clinical conditions underlying body image disturbance. These results may also have implications for future research looking to inform early interventions in body image disturbance, suggesting that addressing impaired processing of affective interoceptive stimuli may be the key to current treatment approaches available not only for AN but also for BDD populations.

## Authors’ contributions

VC conceived the study. SSa contributed to the concept and design of the study. AM and PDT helped with implementation of the paradigm. SSh performed data collection. VC analysed the data, with input from SSa. The first draft of the manuscript was written by VC, with support from all authors. All the authors approved the manuscript before submission.

## Data availability statement

All the relevant data are provided in anonymized datasets within the paper and its Supporting Information files.

